# Thermal stress drives seagrass fragmentation in the Mediterranean Sea

**DOI:** 10.64898/2026.02.23.707362

**Authors:** Àlex Giménez-Romero, Tomàs Sintes, Carlos M. Duarte, Manuel A. Matías

**Author notes:** Corresponding author: Àlex Giménez-Romero.

## Abstract

*Posidonia oceanica* meadows, which underpin Mediterranean coastal ecosystems, are undergoing accelerated decline, partly driven by thermal stress. While previous quantitative studies have identified temperature thresholds beyond which seagrass mortality increases sharply, we show the cumulative and sublethal impacts of prolonged warming under fluctuating subthreshold conditions. To capture these effects, we introduce Stress Degree Days (SDD), a physiologically grounded index derived from an experimentally validated mortality rate function. Using sea surface temperature (SST) data, we quantified the cumulative thermal exposure across the Mediterranean Basin from 2000 to 2020. Leveraging high-resolution satellite imagery and deep learning–based habitat mapping, we linked SDD-derived thermal exposure to meadow fragmentation, which is a proxy for seagrass health. Our results show that high thermal stress (> 50%) is concentrated along the southern and eastern Mediterranean, where meadows exhibit more than 40% cover loss and elevated fragmentation, even though maximum SSTs remained below lethal limits (LT_50_ = 28.9 °C). This finding highlights the critical role of chronic sublethal thermal stress in driving structural degradation. Future projections under the RCP8.5, ‘business as usual,’ and the more moderate RCP4.5 climatic scenarios indicate basin-wide regression, with expected cover losses of approximately 80% and 40%, respectively, by 2100, and near-total habitat suitability collapse in the southern regions. Consequently, fragmentation indices are projected to double or triple, further disrupting clonal connectivity, sediment retention, and oxygen export. In summary, by integrating physiological mechanisms, large-scale remote sensing, and climate modeling, the SDD framework identifies thermal hotspots, reveals emergent vulnerability patterns, and offers a predictive tool to guide conservation strategies in warming oceans.

## Introduction

Seagrass meadows are foundational coastal ecosystems that sustain marine environments across all continents except Antarctica. These ecosystems provide essential services for human health, livelihoods, biodiversity, and climate regulation [1–3]. The dense canopies of seagrass meadows support coastal biodiversity by providing food and habitat for numerous marine species, including commercially valuable ones. Moreover, seagrasses play a crucial role in sediment stabilization and coastal protection by reducing wave and current impacts [4, 5], promoting sediment deposition [6–8], and anchoring the seabed through extensive rhizome networks [9]. Furthermore, seagrasses are among the most effective natural carbon sinks on Earth, sequestering atmospheric CO_2_ in both above- and belowground biomass and in long-lived sediment pools [10–12]. Despite their immense ecological and economic value, seagrass meadows are declining globally, with approximately one-third of the global area lost since World War II [3, 13]. The Mediterranean Basin has been particularly affected by local stressors impacting seagrass meadows, including eutrophication, habitat degradation, and mechanical disturbances, which are further compounded by climate change [14].

The Mediterranean Sea is warming faster than the global ocean average, threatening endemic species such as *Posidonia oceanica*, the region’s dominant seagrass species [15, 16]. Climate projections predict that rising sea surface temperatures (SST), particularly under the RCP8.5 scenario, will drastically reduce suitable habitats for *P. oceanica*, with niche models forecasting losses exceeding 75% by 2050 and potential functional extinction by 2100 [17, 18]. However, these estimates may be conservative, as they rely primarily on the maximum tolerable temperatures (thermal limits) for species survival, thereby neglecting cumulative sublethal thermal stress [19]. Seagrass thermal responses are typically inferred from laboratory experiments that expose plants to prolonged, constant temperatures to assess mortality and physiological performance [20]. While these controlled experiments reveal fundamental thermal tolerance limits, they may not capture the cumulative stress imposed by the dynamic and fluctuating conditions that seagrasses encounter in the wild. In situ, seagrasses experience temporally variable temperature regimes, and mortality is likely the result of integrated stress over time rather than instantaneous threshold exceedance.

Recent advances in artificial intelligence (AI) and machine learning (ML) have enabled seagrass mapping using airborne and satellite imagery [3], yielding promising but geographically constrained results [21–28]. Most existing models are designed for local applications and struggle to generalize across the diverse environmental conditions of the Mediterranean Basin, limiting their scalability for large-scale monitoring. Recently, however, a convolutional neural network model (CAMELE) has been developed to overcome this limitation [29]. CAMELE was trained on a geographically diverse benthic habitat dataset spanning the Balearic Islands, accounting for variations in water depth, water turbidity, and satellite viewing geometry, and was validated across independent regions. It achieved consistent, high-accuracy classification of *Posidonia oceanica* cover at ecologically meaningful spatial scales, with a median performance of 77.30% in regions substantially different from the training domain and up to 95.22% across the entire dataset. This open-source framework enables basin-wide assessments of seagrass distribution and structural changes, providing a baseline for evaluating climate-driven impacts, such as ocean warming and habitat fragmentation.

In this study, we integrated high-resolution remote-sensing-derived habitat maps from over 30 satellite images spanning diverse Mediterranean regions, generated using the CAMELE model, with historical sea surface temperature (SST) time series and experimental physiological data to resolve the impacts of cumulative thermal stress on *P. oceanica* meadows across the Mediterranean Sea. We introduce Stress Degree Days (SDD), a novel metric that quantifies thermal exposure by convolving daily SST data with an experimentally derived mortality rate curve, thereby capturing integrated thermal stress across natural variability. Using AI-driven habitat classifications, we computed seagrass cover and fragmentation across the Mediterranean Sea, revealing structural differences at basin-wide scales. Our results show that thermal stress correlates strongly with reduced cover and increased fragmentation, even though maximum SSTs remained below lethal limits, highlighting how sublethal thermal stress undermines ecosystem resilience. This integrated approach, which combines physiology, remote sensing, and modeling, provides a robust predictive framework for understanding seagrass decline in warming oceans, identifying emergent patterns beyond simplistic thresholds, and guiding targeted conservation initiatives amid escalating biodiversity threats.

## Results

### Quantifying thermal stress in the Mediterranean Sea

We developed a physiological model to estimate the cumulative thermal stress experienced by *Posidonia oceanica* meadows across the Mediterranean Sea. The model integrates daily sea surface temperature (SST) data with an experimentally derived mortality-rate function to compute a time-accumulated index of thermal exposure, which we term *Stress Degree Days* (SDD) (Fig. 1a,b; Methods). Using experimental data from 25-day exposures of *P. oceanica* shoots to constant temperature conditions, the accumulated SDD values were linked to observed shoot mortality through a logistic relationship (Fig. 1c). When applied to spatially and temporally resolved SST data, the resulting quantity—hereafter referred to as thermal stress—is a proxy for the expected shoot mortality associated with cumulative thermal exposure. By integrating both the magnitude and duration of supra-optimal temperatures, this metric captures the progressive physiological burden imposed by warming under naturally fluctuating field conditions. In this sense, the SDD approach is similar to the degree heating weeks and days metric of stress, developed to predict the risk of bleaching and mortality in corals, which provides the basis for many models projecting future coral loss cover under climate change [30].

**Figure 1:**
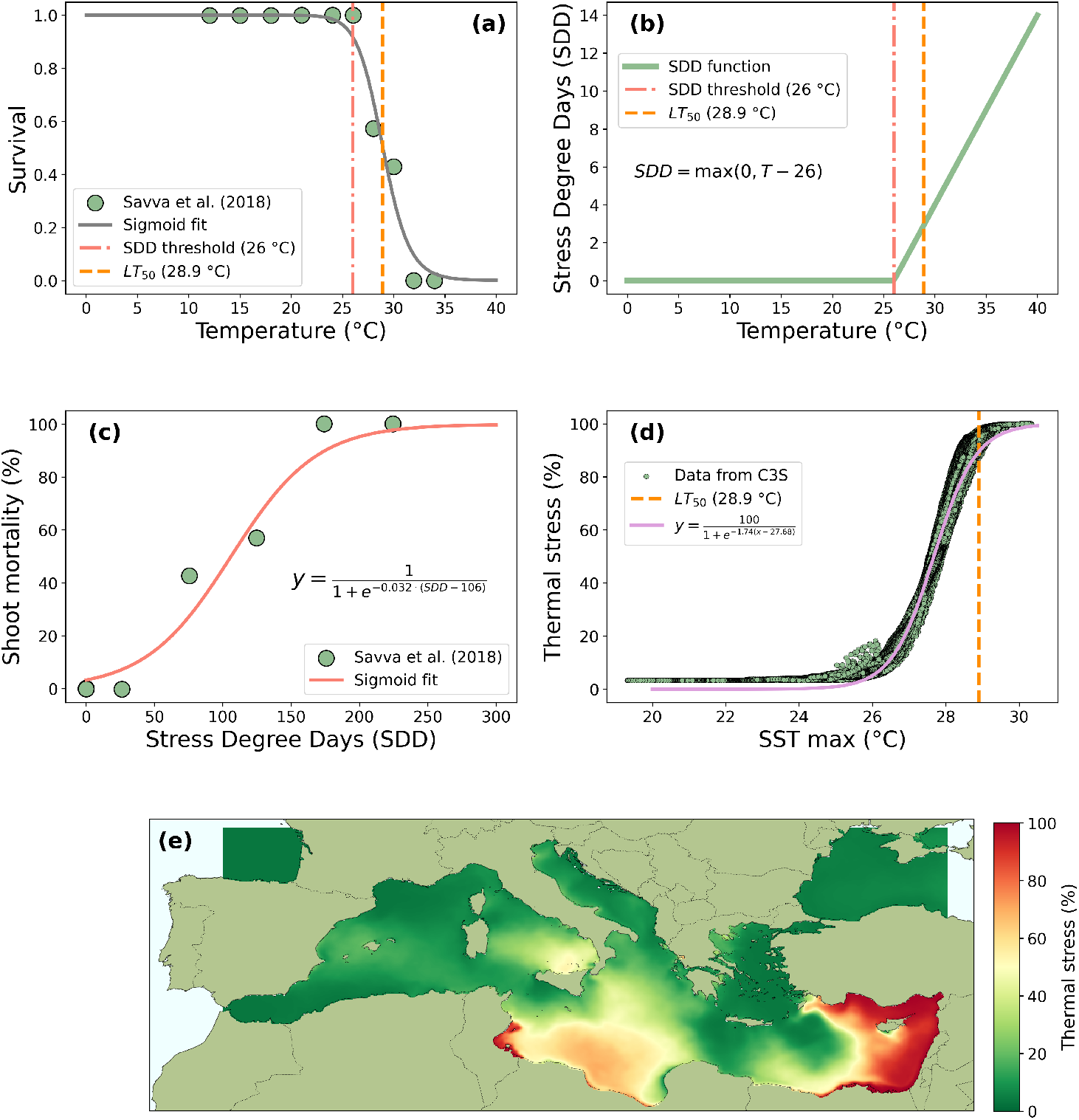
Temperature-dependent mortality model for *Posidonia oceanica*. (a) Experimental survival of *Posidonia oceanica* shoots after 25 days of exposure to constant temperatures, based on data from [20]. The dashed-dotted red line denotes the threshold below which mortality was not observed and is used to compute thermal stress. The orange dashed line, *LT*_50_, corresponds to the lethal temperature threshold employed in previous studies to quantify the mortality risk in *Posidonia oceanica* based on the maximum SST [20]. (b) Stress Degree Days (SDD) as a function of temperature with a threshold of *T* = 26 °C recalculated from the experimental data, which showed negligible mortality below 26 °C and a steep increase at higher temperatures. (c) Logistic fit of experimentally measured shoot mortality after 25 days under constant temperature conditions to SDD, computed from the experimental temperatures. (d) Thermal stress (expressed as predicted shoot mortality) as a function of the maximum SST across a 5 km grid in the Mediterranean Sea (green dots), showing a logistic relationship (pink line). We observe that the commonly used thermal limit *LT*_50_ captures only the highest percentile of thermal stress. (e) Spatial distribution of the average thermal stress in the Mediterranean Sea in the period 2000–2020.

In contrast to traditional threshold-based approaches, which treat thermal stress as a discrete exceedance of fixed temperature limits, our framework explicitly accounts for the cumulative nature of thermal stress over time. The model reveals that the maximum monthly average temperature (SST_max_), a commonly used proxy for thermal stress, exhibits a logistic relationship with the SDDs (Fig. 1d). However, our findings indicate that focusing exclusively on lethal temperature thresholds, such as the experimentally derived *LT*_50_ = 28.9 °C, provides an incomplete view of the stress dynamics. These thresholds capture only the uppermost tail of the thermal stress distribution (above the 90th percentile). In contrast, substantial cumulative stress can still accumulate in regions where SST_max_ remains below the lethal limits. This highlights the importance of accounting for sublethal and time-dependent effects when assessing the vulnerability of seagrass ecosystems to warming.

Using the dependence of thermal stress on SDDs, we generated spatially explicit maps of thermal stress across the entire Mediterranean Basin, utilizing daily SST data from the Copernicus Climate Change Service (C3S) for the period 2000–2020 (see Methods). Our spatial analysis showed that areas of high cumulative thermal stress (i.e., > 50%) are currently concentrated along the southern and eastern coasts of the Mediterranean, particularly off Tunisia, Libya, Egypt, Gaza, Lebanon, Syria, and Turkey (Fig. 1e). According to our analysis, these regions correspond to zones where sustained exposure to elevated temperatures has already exceeded the physiological tolerance limits inferred from experimental data, suggesting that these meadows may approach critical thresholds of decline under future climate conditions.

### The impact of thermal stress in *Posidonia oceanica*

We integrated satellite-derived habitat classifications with the SDD metric generated by our physiological model to examine the relationship between cumulative thermal stress and the spatial structure of *Posidonia oceanica* meadows across the Mediterranean Basin. To this end, for each study site, *P. oceanica* cover was estimated using our deep learning framework [29], which provides spatially consistent predictions of the extent of seagrass from multispectral satellite imagery (see Supplementary Figs. 1 to 3 for examples of habitat mapping). Meadow fragmentation was quantified using two complementary indices that capture distinct aspects of spatial configuration: (i) a gap-weighted fragmentation index (FI) sensitive to the size and shape of internal sand gaps within meadows and (ii) a composite fragmentation index derived from standardized landscape metrics describing patch density, subdivision, and compactness (see Methods). Together, these metrics of fragmentation provide a robust quantification of meadow structures across spatial scales and environmental settings.

Our results revealed a clear correspondence between cumulative thermal stress and seagrass landscape configuration (Fig. 2). The distribution of thermal stress across the Mediterranean Sea (Fig. 1e) mirrors the landscape configuration of the seagrass meadows, with areas of elevated stress concentrated along the southern and eastern coasts, particularly in the Levantine and Tunisian regions (Fig. 2a), which are populated with the lowest values of seagrass cover and the highest degrees of fragmentation (Fig. 2b–c). These results indicate that meadows exposed to sustained sublethal warming exhibit marked structural degradation.

**Figure 2:**
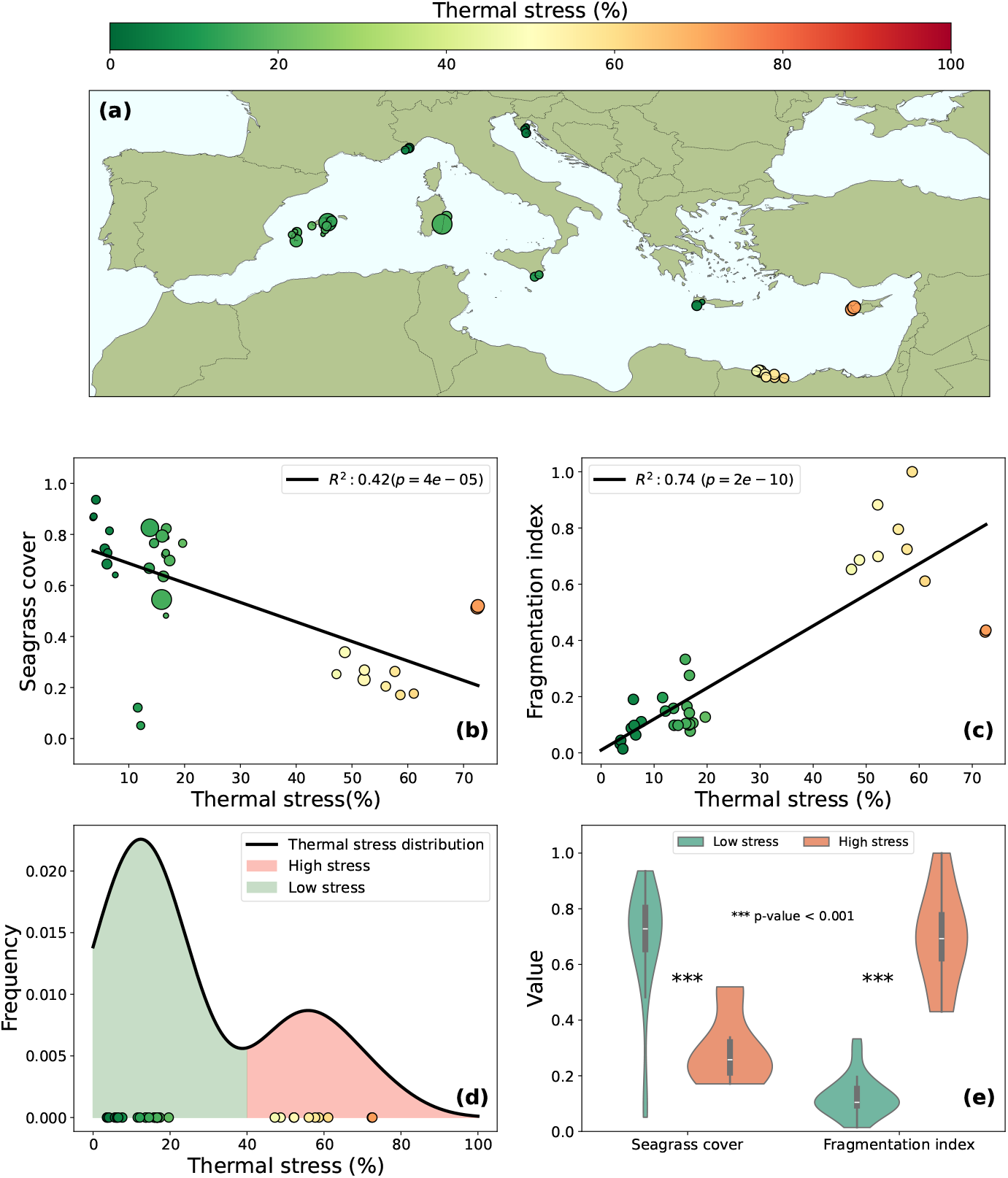
Relationship between sea surface temperature, thermal stress, and *Posidonia oceanica* cover. (a) Locations where satellite imagery was obtained. The color indicates the estimated thermal stress for *Posidonia oceanica* meadows based on the historical SST data. (b) Scatter plot of seagrass cover versus thermal stress, showing a clear negative correlation. (c) Scatterplot of seagrass fragmentation versus thermal stress, showing a clear positive correlation. (d) The distribution of thermal stress across all sites highlights the emergence of two natural stress regimes: low (green) and high (red) stress (separated by a thermal stress level of 40%), which were used to categorize the sampling locations. (e) Violin plots of the distribution of seagrass cover and fragmentation in the two identified regions of thermal stress in our dataset. Regression statistics (*R*^2^, *p*-values) are indicated where applicable.

Categorizing the sites further quantified the relationship between thermal stress and meadow conditions into low- and high-stress regimes based on the bimodal distribution of SDD values (Fig. 2d). Both *P. oceanica* cover and the two fragmentation indices differed significantly between these regimes (Kruskal–Wallis test: *p* < 0.001; Fig. 2e). On average, meadows located in high thermal stress regions displayed a reduction in cover exceeding 40% relative to low thermal stress areas, together with an increase of more than two-fold in fragmentation scores. The two meadow fragmentation metrics yielded consistent results, with both the gap-weighted FI and composite index showing a monotonic increase with cumulative thermal stress (Supplementary Fig. 7).

Notably, these spatial patterns emerged even in locations where the maximum monthly sea surface temperature (SST_max_) remained below the experimentally determined lethal temperature threshold (*LT*_50_) for *P. oceanica* (Supplementary Fig. 5). This suggests that substantial structural deterioration of meadows can occur under persistent sublethal thermal conditions, as captured by the cumulative stress metric. Overall, the basin-wide analysis revealed a clear spatial segregation of healthy and degraded meadows along a thermal stress gradient, supporting the utility of the SDD-based approach for predicting areas where sustained warming has already compromised meadow integrity.

### Projecting seagrass regression under climate change

We then used the validated relationship between meadow landscape configuration indices and thermal stress to forecast the trajectory of *Posidonia oceanica* meadows under projected Mediterranean warming conditions. We computed future thermal stress from projected SST data under two Representative Concentration Pathways (RCPs): RCP4.5, representing an intermediate emissions scenario, and RCP8.5, a high-emissions pathway assuming continued reliance on fossil fuels by the end of the century (“business-as-usual” scenario). These projections, derived from the Copernicus Marine Biogeochemistry dataset [31] (Methods), enabled us to simulate cumulative thermal stress and its cascading effects on seagrass cover and fragmentation from 2006–2100.

Spatially resolved projections for 2100 revealed a pronounced projected escalation in thermal stress across the Mediterranean Basin, with the most severe impacts concentrated in the southern and eastern regions (Fig. 3a). Under RCP4.5, thermal stress values are expected to exceed 80% along the coasts of Tunisia, Libya, Egypt, and the Levantine Sea, encompassing areas that currently support extensive *P. oceanica* meadows. This intensification of thermal stress translates to substantial habitat degradation, with potential reductions in seagrass cover exceeding 60% in these high-stress zones relative to present-day estimates (Fig. 3b). Beyond the expected hotspots of thermal stress in the southern and eastern Mediterranean [17, 32], our projections revealed the development of intense, geographically constrained thermal hotspots and the critical importance of thermal refugia across the basin (Fig. 3a). Even under the less severe RCP4.5 scenario, areas with previously low stress are projected to experience a sharp increase in cumulative thermal stress, potentially leading to meadow degradation. Notably, projections for 2100 show the emergence of new high-stress centers in the northern Mediterranean, particularly around the Balearic Islands and the southwestern coasts of Italy, where the cumulative thermal stress is expected to exceed the 50% threshold by a considerable margin. This northward and eastward expansion of thermal stress dictates the future spatial pattern of meadow degradation. Conversely, certain areas maintained remarkably low SDD values, even under the high-emissions RCP8.5 pathway (Supplementary Figs. 10 and 11). These include the coasts of southern Spain (Alboran Sea), southern France (Gulf of Lion), and parts of the Aegean Sea, suggesting that these regions will function as critical thermal refugia for *P. oceanica*, potentially safeguarding its presence in the Mediterranean. These areas are projected to warm less than other areas because of their distinct oceanographic regimes. The Alboran Sea is a frontal zone where the surface Atlantic and Mediterranean waters converge and often experience intense mixing due to several processes [33], similar to the Gulf of Lion, which is affected by strong mistral winds leading to convective mixing [34], while the Aegean thermal refugia arise due to the interface cooling due to mixing by cyclones [35].

**Figure 3:**
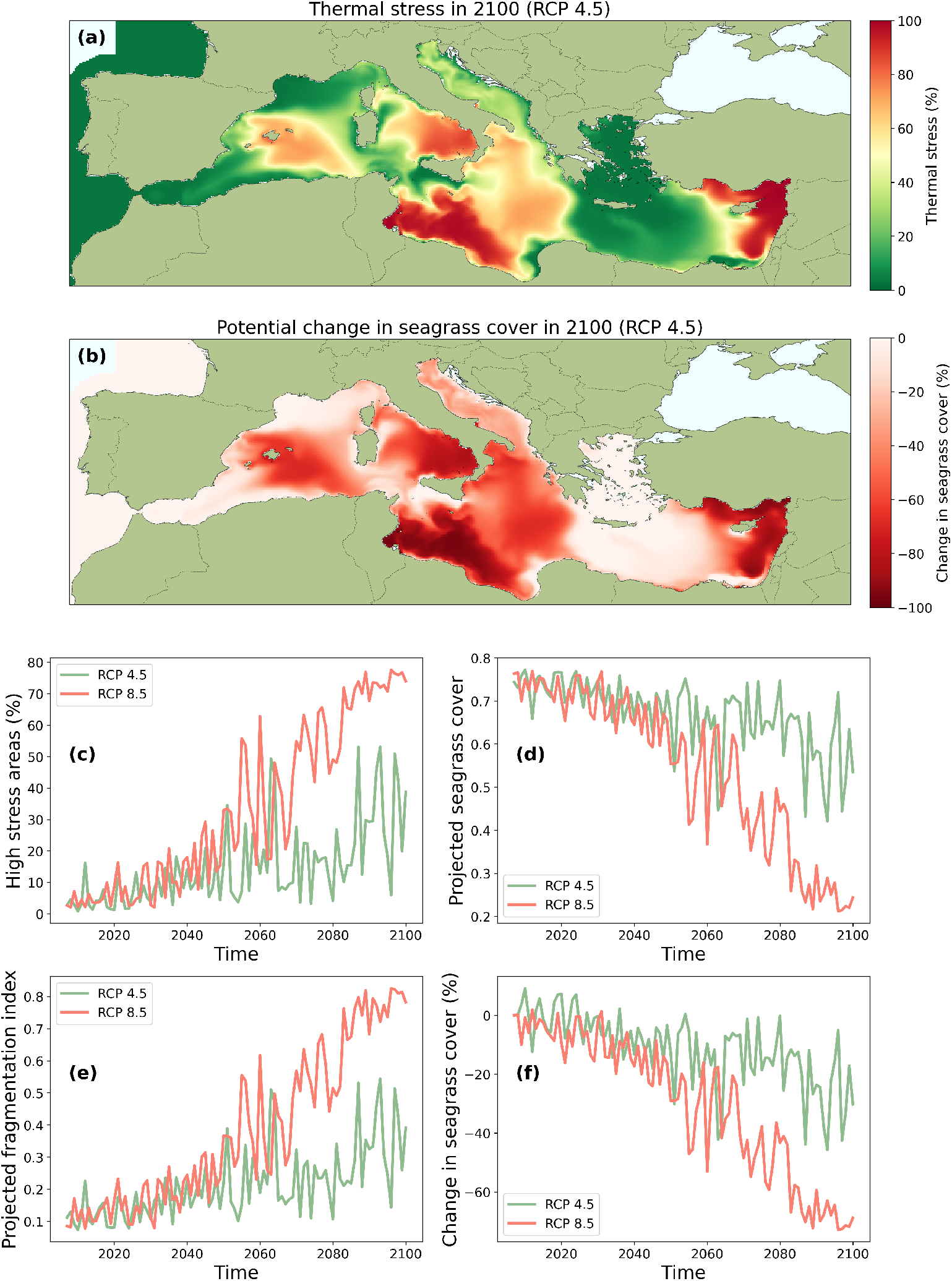
Projected thermal stress and changes in *Posidonia oceanica* meadows under future climate scenarios. (a) Spatial distribution of cumulative thermal stress (SDD) across the Mediterranean Sea in 2100 under RCP4.5. (b) Potential change in seagrass cover (%) in 2100 under RCP4.5, relative to the 2020 baseline estimates. (c) Temporal evolution of the fraction of high-stress areas (> 50%) from 2020 to 2100 under RCP4.5 (green) and RCP8.5 (red). (d) Projected basin-wide seagrass cover from 2020 to 2100 under RCP4.5 (green) and RCP8.5 (red) emission scenarios. (e) Projected fragmentation index from 2020 to 2100 under RCP4.5 (green) and RCP8.5 (red). (f) Cumulative change in seagrass cover (%) from 2020 to 2100 under RCP4.5 (green) and RCP8.5 (red). Projections are based on SST data from the Copernicus Marine Biogeochemistry dataset [31], integrated with the SDD framework.

The empirical link between high cumulative stress and meadow structural deterioration suggests that the extensive areas projected to experience SDD values above 70% will undergo a widespread transition from continuous, stable meadows to highly fragmented landscapes (Supplementary Fig. 12), which is an unstable configuration that precedes ecosystem collapse [36]. This transition, characterized by the breakdown of meadow connectivity and loss of essential habitat functions, is particularly acute in current and emerging thermal hotspots. By 2100, under the high-emissions scenario, this predicted structural collapse will transform the vast majority of the Mediterranean basin from a resilient seagrass ecosystem into an environmentally simplified, low-cover, high-fragmentation seascape, with the last remnants of continuous cover relegated to the identified northern thermal refugia (Supplementary Figs. 10 to 12).

Temporal dynamics underscore progressive deterioration, with the fraction of high-stress areas (> 50%) expanding nonlinearly over the next few decades (Fig. 3c). Under RCP4.5, this fraction is projected to increase from approximately 10% in 2020 to over 40% by 2100, whereas RCP8.5 accelerates this trend, potentially affecting more than 50% of the basin by mid-century and approaching near-ubiquitous stress (> 70%) by 2100. Concomitantly, basin-wide seagrass cover is expected to decline steadily, with RCP4.5 yielding a 30-40% reduction and RCP8.5 precipitating losses exceeding 70% by the end of the century (Fig. 3d,f). Fragmentation indices mirror this regression, exhibiting a twofold to threefold increase under RCP4.5 and even steeper amplification under RCP8.5, indicative of accelerating meadow disassembly into isolated patches (Fig. 3e).

The preceding analysis focused on basin-scale average impacts; however, it is critical to recognize that thermal stress is inherently depth-dependent. As *P. oceanica* meadows can extend down to 50 m, we investigated how the projected cumulative thermal stress varies with depth under the RCP4.5 scenario (Fig. 4). Our results confirm that the shallowest depths (e.g., < 10 m) are projected to experience significantly greater thermal stress than deeper zones (e.g., > 20 m) by 2100 (Fig. 4 (a),(b)). While thermal stress values at 10 m frequently exceeded 70% across large areas, they were markedly attenuated at 20 m. This differential warming is quantified by the percentage of the Mediterranean seabed area falling into the high-stress category (> 50%) as a function of depth (Fig. 4 (c)). This analysis revealed a profound increase over time in the extent of high-stress zones, but with a sharp decline in vulnerability beyond approximately 20 m depth. For the period 2090-2100, the extent of the high-stress area is nearly four times greater at 10 m than at 30 m.

**Figure 4:**
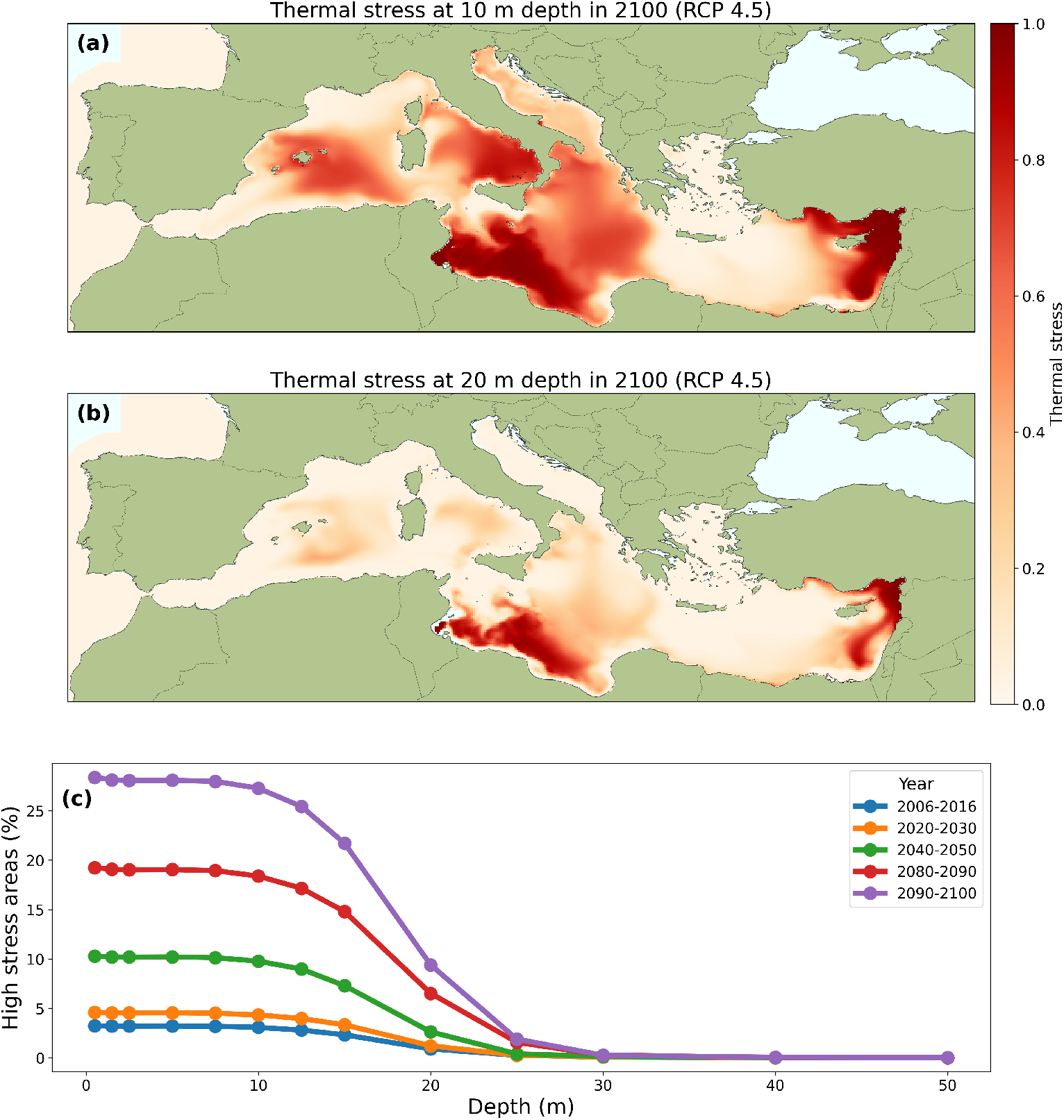
Projected thermal stress across depths under the RCP4.5 scenario. (a) Spatial distribution of cumulative thermal stress (SDD) at 10 m depth in 2100 under the RCP4.5 scenario. The redder areas indicate higher thermal stress, ranging from 0 (no stress) to 1 (maximum stress). (b) Spatial distribution of cumulative thermal stress (SDD) at 20 m depth in 2100 under the RCP4.5 scenario, showing a reduced intensity compared to 10 m. (c) Average percentage of high-stress areas (> 50%) across the Mediterranean Sea as a function of depth for different time periods (2006-2016, 2020-2030, 2040-2050, 2080-2090, and 2090-2100). The plot illustrates a projected increase in high-stress areas over time, with a sharp decline in their extent beyond 20-30 m depth, highlighting the shallow-water vulnerability.

## Discussion

Our findings show that cumulative thermal stress, quantified using the Stress Degree Days (SDD) metric, is a robust predictor of *Posidonia oceanica* meadow degradation across the Mediterranean Sea. By integrating experimental mortality data with temporally varying sea surface temperatures, the SDD approach effectively captures sublethal thermal stress, which is overlooked by traditional absolute-threshold-based metrics. This distinction is critical in naturally fluctuating marine environments, where prolonged exposure to moderately elevated temperatures induces progressive physiological damage, manifested as reduced shoot density, impaired growth, and eventual meadow fragmentation. Recent empirical evidence supports this hypothesis. Marine heatwaves (MHWs) exacerbated by climate change impose cumulative sublethal stress on *P. oceanica*, triggering stress-induced adaptive responses such as profuse flowering, a stress-induced reproductive strategy [37, 38]. Moreover, ontogeny-specific vulnerabilities reveal that seeds exhibit heightened sensitivity to thermal events [39], reinforcing the importance of cumulative metrics, such as SDD, that integrate impacts across developmental stages. Consequently, our approach extends beyond previous threshold and thermal niche models, which likely underestimate the contribution of sublethal thermal stress to species decline [40, 41].

The spatial patterns revealed in our analysis correspond closely with the documented declines in *P. oceanica* meadows in the southern and eastern Mediterranean [32], where warming rates are increasing most rapidly. Regions such as Tunisia, Libya, and the Levantine coast exhibit high SDD values and a prevalence of fragmented meadows with over 40% loss of cover, even where maximum temperatures remain below lethal limits. This indicates that chronic cumulative stress progressively undermines meadow resilience through processes such as inhibited clonal expansion and heightened vulnerability to feedback mechanisms [36, 42]. Transcriptomic analyses of simulated MHWs further revealed differential gene expression across stress intensities [43], reinforcing the notion that prolonged sublethal stress drives structural degradation. The ecological consequences of fragmentation extend beyond structural decline and affect ecosystem metabolism and resilience. Recent field evidence from Mediterranean *P. oceanica* meadows shows that moderate fragmentation (FI > 3.4), combined with enhanced hydrodynamic flow, significantly increases dissolved oxygen concentrations through improved lateral advection and reduced boundary layer thickness, thereby enhancing both gross primary production and net community oxygenation [44]. This suggests a counterintuitive adaptive mechanism: while initial fragmentation accelerates clonal disconnection, it also boosts local productivity and oxygen export, which may support neighboring benthic communities and delay hypoxia-driven feedback in warming coastal embayments. However, this metabolic advantage is likely transient; as thermal stress intensifies and fragmentation exceeds critical thresholds, declining canopy complexity and rhizome connectivity ultimately impair carbon sequestration, sediment stabilization, and long-term resilience.

Projections under the RCP4.5 and RCP8.5 scenarios predict severe habitat regression, with potential cover loss exceeding 70% under high-emission conditions. Such declines would amplify habitat fragmentation and increase the risk of functional extinction in the southern basins [16, 17]. These projections align with broader trends across Mediterranean ecosystems, where synergistic stressors, including sea-level rise, invasive species, and marine heatwaves, trigger cascading biodiversity losses and diminished carbon sequestration [45]. Sustained warming is expected to simplify meadow architecture progressively, thereby disrupting clonal connectivity and rhizome-mediated resource sharing. This breakdown can accelerate local extinction dynamics while eroding the genetic diversity essential for long-term adaptation [46]. As canopy density declines, sediment retention capacity weakens, increasing erosion and turbidity and inhibiting recolonization, thereby reinforcing feedback loops that strengthen habitat degradation. In semi-enclosed coastal systems, these structural losses diminish the role of meadows as metabolic hotspots, reducing oxygen export to adjacent habitats and weakening support for associated fauna. The convergence of thermal stress with co-occurring pressures ultimately risks pushing *P. oceanica* meadows beyond critical tipping points, fragmenting once-continuous seascapes into lower-functioning patches with diminished ecological roles [42].

Although our model represents a significant advancement in quantifying cumulative thermal stress on *Posidonia oceanica* meadows by integrating experimental mortality data with satellite-derived sea surface temperature (SST) records, several limitations must be acknowledged to contextualize its applicability and guide future refinements. First, reliance on SST as a thermal proxy assumes uniform heat transfer to benthic habitats; however, this approach overlooks vertical thermal stratification. Deeper meadows, which often extend to 40 m or beyond [47], may experience substantially cooler and more stable conditions than surface waters, potentially leading our model to underestimate the resilience of bathymetrically diverse sites. Our depth-resolved model reveals depth-dependent vulnerability, with shallow (< 10 m) meadows likely to experience greater thermal stress and landscape fragmentation than deeper ones. Hence, our results suggest that shallow *P oceanica* meadows are critically threatened by climate change and are likely to be the first to undergo structural collapse. Deeper meadows may provide refuge within particular regions, offering a narrow but increasingly important window of persistence for the species. However, shallow, < 10 m, *P. oceanica* meadows play a disproportional role in coastal protection [48]. Accordingly, whereas *P. oceanica* will continue to support biodiversity, its coastal protection benefits will be severely impaired.

The model was based on experimental data from the western Mediterranean (Balearic Islands) and may not necessarily apply to the entire range of *P. oceanica*. Unfortunately, the lack of controlled thermal experiments testing mortality responses to increasing temperatures precludes the testing of these models elsewhere. The existence of a thermal gradient reflected in increasing maximum temperatures from the western towards the eastern Mediterranean basin could imply the a priori expectations of eastern Mediterranean *P. oceanica* meadows are more resistant to thermal stress than those in the western Mediterranean. However, existing evidence does not suggest the existence of such variability. For instance, reconstruction of growth dynamics in Greek *P. oceanica* populations concluded that maximum temperatures above 26.5 °C caused a sharp decrease in production [32], indicating a threshold for sublethal thermal stress similar to that empirically resolved for western Mediterranean populations [20]. Likewise, reciprocal transplant experiments across the western, central, and eastern Mediterranean [49] and across cool subthermocline environments to warmer, shallower environments [50] did not show any correspondence between the thermal origin of the transplants and their performance. Hence, existing evidence does not reject the assumption of a consistent thermal response of *P. oceanica* across its entire distribution range.

Another limitation of our framework is that it does not explicitly account for the multi-stressor interactions prevalent in coastal ecosystems. Synergistic effects between warming and eutrophication can amplify nutrient toxicity and reduce plant performance under chronic co-exposure [51, 52]. Similarly, herbivory may intensify under elevated temperatures, potentially exacerbating fragmentation through selective grazing [53], while ocean acidification coupled with marine heatwaves (MHWs) impairs photosynthetic efficiency and seedling survival [54, 55]. These interactions could substantially modulate SDD estimates but remain unmodeled in the current framework, as they require local level knowledge of current and future stressors, which is lacking. However, increased awareness of the importance of *P. oceanica* is driving conservation efforts, particularly on the northern coast of the Mediterranean Basin [56]. Enhanced conservation management should reduce the local pressures exerted on these ecosystems, further supporting increased restoration efforts [57].

The precision of the model reported here is further limited by constraints imposed by remote sensing. Deep-learning-based habitat mapping, while robust across heterogeneous conditions, is constrained by satellite imagery limitations, including atmospheric interference, water turbidity, and spectral confusion with other benthic organism factors that can reduce classification accuracy to below 80% in turbid or deep sites [29]. Moreover, the 3–4 m resolution of PlanetScope data may fail to resolve fine-scale fragmentation patterns such as small canopy gaps or patchy die-offs, while fragmentation metrics are sensitive to the spatial scale at which they are calculated. Temporal mismatches between historical SST records (2000–2020) and recent imagery (2023) introduce additional uncertainty when linking current meadow states to past thermal stress, potentially overlooking recovery dynamics and lagged mortality responses. Finally, future projections under RCP4.5 and RCP8.5 inherit uncertainties from climate models [58]. The assumptions of static species distributions ignore potential range shifts, selection, and adaptive evolution [59], which may shift thermal sensitivity curves under future climate change, as reported for corals [30]. This could lead to an overestimation of the decline in resilient populations or an underestimation of compounding factors, such as invasive species, that accelerate degradation.

More generally, a key limitation of our study is its focus on locations selected as relatively pristine, under the assumption that fragmentation in these areas is predominantly driven by thermal stress rather than by direct human disturbance. This strategy allowed us to isolate the effects of cumulative sublethal warming on meadow structure; however, it also introduces uncertainty. Some sites may not be as undisturbed as presumed, and low-intensity or cryptic anthropogenic pressures—such as occasional anchoring, historical degradation, or diffuse coastal impacts—may still contribute to fragmentation patterns in ways not captured by our screening criteria. Consequently, while our results strongly support a link between cumulative thermal stress and spatial degradation, the influence of undetected local disturbances cannot be ruled out entirely. In addition, this focus on minimally impacted sites implies that our estimates represent a conservative lower bound for the basin-wide drivers of degradation. In more heavily affected areas, where thermal stress interacts synergistically with local anthropogenic pressures, fragmentation rates are likely to be considerably higher, and ecological thresholds are likely to be reached more rapidly.

Addressing these limitations through integrated approaches, including in situ validation, multi-depth thermal profiling, coupled multi-stressor experiments, and the deployment of higher-resolution sensors, will substantially enhance the predictive accuracy. Likewise, future applications of this framework should explicitly integrate quantified gradients of human impact to disentangle better the interacting roles of warming and local stressors in seagrass fragmentation. Overall, we expect our modeling framework to inform adaptive management strategies for conserving this keystone species under accelerating climate change.

## Methods

### Temperature data

We used sea surface temperature (SST) data from two complementary sources to quantify thermal exposure in the *Posidonia oceanica* meadows. For the historical analysis, we employed the dataset “Sea surface temperature daily data from 1981 to present derived from satellite observations” [60] provided by Copernicus Climate Change Service (C3S). This product provides daily global SST estimates derived from multiple infrared satellite sensors and processed using optimal interpolation techniques. The dataset merges multi-sensor inputs to produce spatially complete gap-free temperature fields on a regular 0.05° grid (∼ 5 km resolution), ensuring consistent and homogeneous coverage across space and time.

For future climate projections, we used temperature data from the Copernicus dataset “Marine biogeochemistry data for the Northwest European Shelf and Mediterranean Sea from 2006 to 2100 derived from climate projections” [31]. This dataset is based on simulations produced using the European Regional Seas Ecosystem Model (ERSEM v15.06) coupled with the regional ocean circulation models POLCOMS and NEMO through the FABM framework. The models account for physical processes (e.g., temperature, salinity, and currents) and biogeochemical cycles (e.g., nutrients and oxygen) and are forced by downscaled CMIP5 climate projections using the RCA4 atmospheric model. SST was extracted under two Representative Concentration Pathways (RCPs): RCP4.5 and RCP8.5. These projections were used to estimate future thermal stress values and assess potential impacts on seagrass fragmentation under climate change scenarios. The dataset provides daily temperature data on a regular 0.1° latitude–longitude grid (∼10 km resolution) over the Mediterranean Basin for the period 2006-2100 at different depths. For consistency with the previous data source, we used temperature data from the first depth level of *z* = 0.5 m.

### Modelling seagrass thermal stress

To model the impact of thermal stress on *P. oceanica* cover, we used the experimental results from Savva et al. [20], which quantified shoot mortality as a function of prolonged exposure to a constant temperature. Specifically, the authors reported the percentage of shoot mortality after 25 days of exposure to a constant temperature at multiple thermal levels. Thus, the results of the experiment represent the cumulative effect of a temperature-dependent mortality rateµ(*T*), which is constant over time in the experimental setup. However, seagrasses are exposed to time-varying temperatures throughout their lifetime; therefore, a model that accounts for the dynamic nature of thermal stress is necessary.

We modeled the thermal stress as the cumulative effect of a temperature-dependent mortality rateµ(*T* (*t*). We consider that, under stress, the shoot density of a seagrass population follows the dynamics

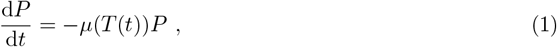

so that the shoot density at time *t* is obtained by solving the differential equation

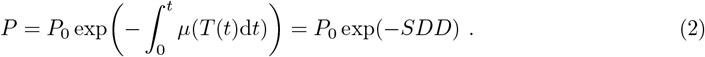

Thus, Eq. (2) models the potential exponential decrease in *Posidonia oceanica* shoot density as a function of cumulative temperature-dependent mortality 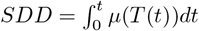, hereafter called *Stress Degree Days*.

The temperature-dependent mortality rateµ(*T*) was extrapolated from the experimental results of Savva et al. [20] by considering no stress below a certain threshold and a linearly increasing mortality rate with temperature above the threshold. Altogether, this is modeled using a nonlinear piecewise function,

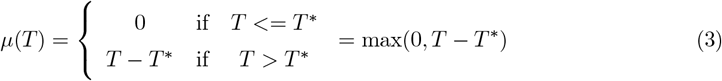

The value of the threshold, *T* ^*^, corresponds to the lowest temperature at which there is no mortality in the experiments of Savva et al. [20], that is, *T* = 26 °C (Fig. 1 (a) and (b)).

Finally, the accumulation of SDD was linked to the thermal stress by fitting a logistic curve to the experimental results of Savva et al. [20] (Fig. 1 (c)). Because all experiments were performed for the same amount of time, Δ*t* = 25 days, and at different **constant** temperatures, the SDD of each experiment is easily computed as,

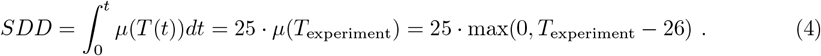

Finally, the relationship between SDD and experimentally-derived mortality is described by a logistic function, which allows the translation of the accumulated stress into an estimate of potential mortality (Supplementary Fig. 4), thereby named “thermal stress”,

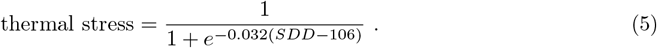

### Thermal stress from SST data

In practice, the integral of the time-varying mortality rate is obtained computationally from temperature data by approximating the integral by a summation,

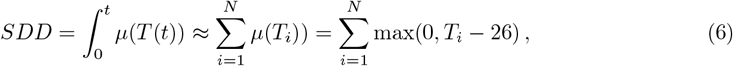

where *T*_*i*_ corresponds to the daily SST values.

### Bathymetry data

Bathymetry data were obtained from the European Marine Observation and Data Network (EMODNET) [61]. EMODnet Bathymetry provides a service for viewing and downloading a harmonized Digital Terrain Model (DTM) for European sea regions. The data consisted of GeoTIFF layers with ∼ 100 m pixel resolution of mean depth values, which were interpolated to the satellite imagery resolution.

### Satellite data

Satellite imagery was obtained from Planet under the Education and Research Program, which provides limited, noncommercial access to PlanetScope and RapidEye imagery [62]. In particular, we acquired PlanetScope images from the Super Dove (PSB.SD) instrument, which includes Coastal Blue, Blue, Green I, Green, Yellow, Red, Red Edge, and NIR, beginning in 2020. Surface Reflectance (SR) products were selected to ensure consistency across varying atmospheric conditions, thereby minimizing uncertainty in the spectral response across time and location. Thus, these multispectral imagery products are specially designed for temporal analysis and monitoring applications. SR is derived from the standard Analytic Product (radiance), which is processed to top-of-atmosphere reflectance and then atmospherically corrected to bottom-of-atmosphere reflectance.

We acquired 33 satellite images covering 1233 km^2^ of Mediterranean coastal areas and islands for 2023 (see Supplementary Information, Supplementary Table 1). Images were acquired over several days under clear-sky conditions between June and September, when seagrass and algal biomass were abundant. The chosen sites were those with the least possible human disturbance so that their fragmentation was mainly due to warming rather than human activity.

### Data pre-processing

The NIR band, which is strongly attenuated by water, was used to mask land pixels using a simple clustering algorithm (K-means) and was subsequently replaced with bathymetry data. We then masked all pixels with depths greater than 30 m, as the AI model’s performance is significantly reduced beyond this threshold [29]. The resulting processed satellite images were used as input data to obtain habitat estimates.

### Habitat estimates

We employed the deep learning framework developed by [29] to estimate the spatial cover of different benthic habitats, including *Posidonia oceanica*, across 33 selected coastal locations in the Mediterranean Sea. The framework, based on convolutional neural networks (CNNs), classifies each pixel of the pre-processed satellite imagery into one of four ecologically meaningful benthic classes: (1) rocks and brown algae, (2) sandy bottoms, (3) *Posidonia oceanica*, and (4) other green plants (e.g., *Cymodocea nodosa*). The model was trained on an extensive, georeferenced dataset of benthic habitat maps spanning diverse environmental conditions across the Balearic Sea, achieving robust generalization with a mean Intersection over Union (IoU) of 77.30% for the *P. oceanica* class in independent cross-regional validation and a median IoU of 95.22% in the final consensus model trained on the full dataset [29].

We applied the final, fully trained model to the 33 Planet surface reflectance images. Each image was processed at a native 3–4 m resolution, yielding high-fidelity habitat maps that resolve fine-scale meadow boundaries, internal patch structures, and fragmentation patterns (see Supplementary Figs. 1 to 3 for some examples). The resulting habitat maps, covering a cumulative coastal area of 1233 km^2^, served as the base dataset for all subsequent analyses.

### Estimation of *Posidonia oceanica* cover

The cover of *Posidonia oceanica* in each satellite image was computed as

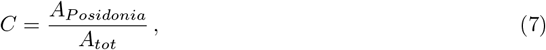

where *A*_*P*_ represents the area of *Posidonia oceanica*, and *A*_*tot*_ is the total area of the studied zone, that is, the area of the satellite image that is classified as any of the benthic categories.

### Fragmentation analysis

To quantify the spatial fragmentation of *Posidonia oceanica* meadows, we computed two complementary fragmentation indices designed to capture distinct ecological aspects of the seagrass spatial structure.

We first calculated a gap-weighted fragmentation index (FI) adapted from the urban fragmentation metric [63, 64] and applied it to seagrass systems to assess the ecological impact of discontinuities on productivity [65]. This index accounts for both the area of bare sand within a defined landscape unit and the spatial configuration of gaps within the seagrass matrix.

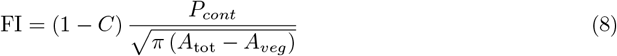

Here, *A*_tot_ is the total area considered, *A*_veg_ is the vegetated area, and *P*_cont_ represents the contact perimeter between the vegetated and unvegetated areas. The first term captures the proportion of the landscape that is unvegetated, whereas the second term describes the geometric complexity of the gaps, weighted by their perimeter-to-area relationship. This index ranges from 0 (fully continuous meadow) to 1 (highly fragmented meadow) and is sensitive to both the abundance and spatial configuration of non-vegetated patches embedded within the meadow.

To complement this measure, we computed a composite fragmentation index based on a suite of landscape-level metrics widely used in landscape and seascape ecology [66–68]. These metrics were calculated from binary habitat maps using the `landscapemetrics` package in R [69] and included the following:

- **Patch density (PD)** – number of discrete seagrass patches per unit area, reflecting the degree of habitat subdivision.
- **Landscape division index (LD)** – the probability that two randomly chosen locations in the landscape belong to different patches, increasing with fragmentation.
- **Area-weighted mean perimeter-to-area ratio (AWMPAR)** – a measure of patch compactness and edge complexity.
- **Mean radius of gyration (GYRATE)** – describes the average distance between cells within a patch, serving as a proxy for patch cohesion.
- **Number of holes** – quantifies internal perforations within seagrass patches and is calculated using the ‘Union’ tool in ArcGIS. This value was normalized per hectare of seagrass to account for variations in overall coverage.

Each metric captures a different dimension of fragmentation: extent, configuration, and internal structure. All metrics were standardized (min-max normalization) and integrated into a single composite fragmentation index [68, 70]. The resulting index ranged from 0 (low fragmentation) to 1 (high fragmentation), providing a unified metric for cross-site comparisons.

Together, these two approaches provide complementary perspectives on seagrass fragmentation: one emphasizes the influence of internal bare-sand gaps, whereas the other captures broader changes in patch configuration and integrity. We used both indices to assess the relationship between fragmentation and thermal stress across the study sites.

## Data and code availability

The CAMELE deep learning model used to estimate *Posidonia oceanica* cover is openly available at [71]. Because satellite imagery was obtained from Planet under the Education and Research Program, which provides limited non-commercial access to PlanetScope and RapidEye imagery, we cannot share the raw images. However, all scripts and workflows used to compute Stress Degree Days (SDD), thermal stress, fragmentation indices, and seagrass cover are publicly available in a GitHub repository [72] and archived in Zenodo [73]. All datasets generated in this study, including the predicted *P. oceanica* habitat maps for the selected regions, SDD and thermal stress values for the Mediterranean Sea (historical period 2000–2020), and projected values under RCP4.5 and RCP8.5 scenarios (2006–2100), are available through the Zenodo repository.

## Acknowledgements

We acknowledge Nuria Marbà, Elvira Mayol, and Damià Gomila for their valuable comments and suggestions. We also acknowledge Planet Labs for access to satellite imagery data under their Education and Research Program. This work was supported by grants TED2021-131836B-I00 (SEDIMENT) funded by the Spanish Ministry of Science and Innovation MICIU/AEI/10.13039/501100011033 and by the European Union NextGenerationEU/PRTR Program; PID2021-123723OB-C22 (CYCLE) and PID2024-156062OB-I00 (CHANGE-ME) funded by MICIU/AEI/10.13039/501100011033 and by ERDF, EU; and CEX2021-001164-M (María de Maeztu Program for Units of Excellence in R&D) funded by MICIU/AEI/10.13039/501100011033. A.G.R. acknowledges financial support from grant JDC2024-053275-I, funded by MICIU/AEI/10.13039/501100011033 and FSE+.

## Author contributions statement

A.G.R., T.S., and M.A.M. conceptualized the study. A.G.R. developed the methodology, performed the formal analysis, curated the data, and created the visualizations. A.G.R. designed and implemented the software for the remote sensing analysis and SDD metric computation. M.A.M., C.M.D., and T.S. supervised the project. M.A.M. and T.S. provided funding. A.G.R. wrote the original draft with contributions from T.S., C.M.D., and M.A.M. All authors reviewed and edited the manuscript and approved the final version.

## Competing interests

The authors declare no conflicts of interest.

## Notes

https://zenodo.org/records/17476416

https://github.com/agimenezromero/Thermal-stress-drives-seagrass-fragmentation-in-the-Mediterranean-Sea/tree/main

### Competing Interest Statement

The authors have declared no competing interest.

