## Supplementary Information for "Thermal stress drives seagrass fragmentation in the Mediterranean Sea"

#### 1 Satellite imagery

We compiled high-resolution satellite imagery to estimate the *Posidonia oceanica* cover across the Mediterranean Basin. The dataset consists of 33 PlanetScope scenes acquired between June and September 2023, corresponding to the peak biomass period. Each scene covered an area of approximately 50–100 $km^2$ and was selected under clear sky conditions to minimize atmospheric distortion. All images were processed for surface reflectance and co-registered to a uniform spatial resolution of 3m. Table 1 lists the acquisition dates and geographic coordinates for each scene.

**Supplementary Table 1. Satellite images used for habitat classification.** A summary of the PlanetScope satellite imagery dataset used to estimate *Posidonia oceanica* cover across the Mediterranean Sea. The table reports the geographic coordinates (to the 4th decimal) and acquisition dates for each site. All images correspond to the summer months (June–September), when seagrass biomass is highest.

| Region | Date | N | W | S | E |
|---|---|---|---|---|---|
| Mallorca | 15 July 2023 | 39.8937 | 3.1142 | 39.7250 | 3.3506 |
| Alexandria | 27 July 2023 | 31.1052 | 28.0292 | 31.0635 | 28.2285 |
| Alexandria | 20 July 2023 | 31.1808 | 27.8779 | 31.0842 | 28.0763 |
| Alexandria | 18 July 2023 | 31.3858 | 27.2432 | 31.2124 | 27.4550 |
| Alexandria | 20 July 2023 | 31.0675 | 28.5062 | 31.0005 | 28.6907 |
| Alexandria | 20 July 2023 | 31.3997 | 27.0704 | 31.3564 | 27.2697 |
| Alexandria | 20 July 2023 | 31.3851 | 27.3152 | 31.2037 | 27.5072 |
| Alexandria | 15 July 2023 | 31.2576 | 27.8013 | 31.0873 | 28.0020 |
| Alexandria | 31 July 2023 | 31.4535 | 26.9801 | 31.3731 | 27.1849 |
| Cabrera | 14 July 2023 | 39.2107 | 2.9445 | 39.1219 | 2.9881 |
| Mallorca | 14 July 2024 | 39.3168 | 3.0098 | 39.2516 | 3.1304 |
| Mallorca | 31 July 2023 | 39.8040 | 3.3176 | 39.6244 | 3.4857 |
| Sardinia | 05 July 2023 | 40.0497 | 9.6796 | 39.8586 | 9.7428 |
| Sardinia | 04 July 2023 | 39.9085 | 9.5061 | 39.0957 | 9.6993 |
| Croatia | 08 July 2023 | 44.9574 | 14.4444 | 44.7694 | 14.4812 |
| Croatia | 17 July 2023 | 44.9357 | 14.2823 | 44.6921 | 14.4010 |
| Croatia | 17 July 2023 | 44.7081 | 14.3210 | 44.5465 | 14.3952 |
| Cyprus | 16 July 2023 | 35.0231 | 32.2727 | 34.7012 | 32.4808 |
| Cyprus | 16 July 2023 | 35.0249 | 32.2769 | 34.7005 | 32.4804 |
| Ibiza | 31 August 2023 | 39.0969 | 1.4476 | 38.9027 | 1.6280 |
| Formentera | 03 August 2023 | 38.7838 | 1.3879 | 38.6634 | 1.5942 |
| Formentera | 03 August 2023 | 38.7427 | 1.3642 | 38.6275 | 1.5951 |
| Greece | 02 August 2023 | 35.2353 | 23.9803 | 35.1803 | 24.1426 |
| Greece | 22 July 2023 | 35.2701 | 23.5339 | 35.2034 | 24.0129 |
| Ibiza | 27 June 2023 | 39.1284 | 1.2782 | 39.0077 | 1.6076 |
| Ibiza | 14 June 2023 | 39.0213 | 1.1828 | 38.8524 | 1.3112 |
| Mallorca | 14 July 2023 | 39.6004 | 2.3327 | 39.4913 | 2.4810 |
| Mallorca | 12 August 2023 | 39.6383 | 3.3230 | 39.5057 | 3.4145 |
| Sicily | 10 August 2023 | 36.8067 | 14.4804 | 36.7278 | 14.6728 |
| Sicily | 05 July 2023 | 36.7333 | 14.8439 | 36.6822 | 15.0139 |
| France | 11 August 2023 | 43.8504 | 7.7284 | 43.7826 | 7.9594 |
| France | 19 August 2023 | 43.8086 | 7.5446 | 43.7618 | 7.7612 |
| France | 15 June 2023 | 43.7854 | 7.3282 | 43.6896 | 7.5386 |

#### 2 Habitat predictions

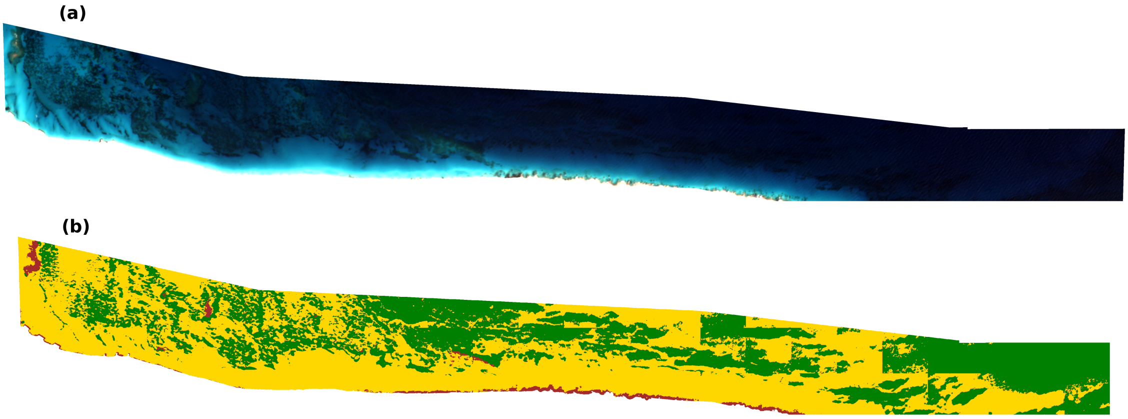


**Supplementary Fig. 1. Example of deep learning-based benthic habitat classification for *Posidonia oceanica*.** (a) True-color composite of high-resolution PlanetScope satellite imagery (3–4 m resolution) for the coast of Alexandria. (b) Corresponding CNN model prediction showing classified habitats: *P. oceanica* (green), bare sediment (yellow), macroalgae/rock (brown), and other green plants (cyan).

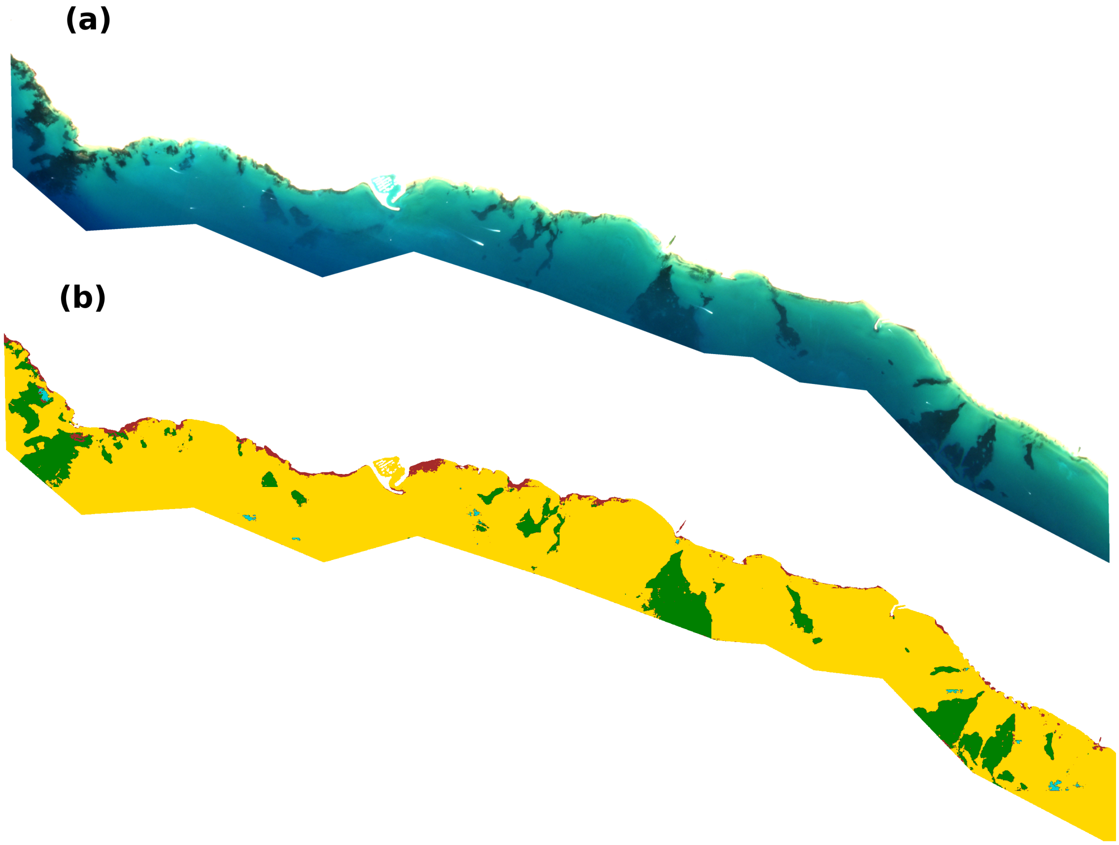


**Supplementary Fig. 2.** (a) True-color composite of high-resolution PlanetScope satellite imagery (3–4 m resolution) for the coast of Sicily. (b) Corresponding CNN model prediction showing classified habitats: *P. oceanica* (green), bare sediment (yellow), macroalgae/rock (brown), and other green plants (cyan).

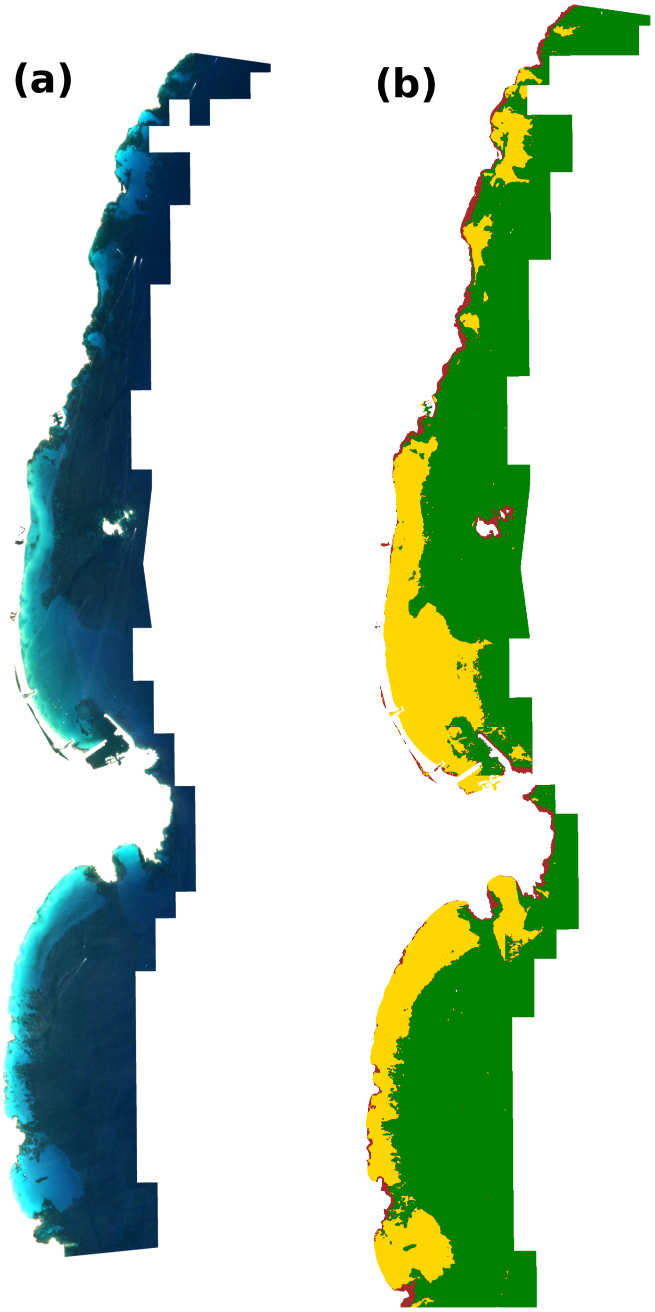


**Supplementary Fig. 3.** (a) True-color composite of high-resolution PlanetScope satellite imagery (3–4 m resolution) for the coast of Sardinia. (b) Corresponding CNN model prediction showing classified habitats: *P. oceanica* (green), bare sediment (yellow), macroalgae/rock (brown), and other green plants (cyan).

#### 3 Modelling thermal stress

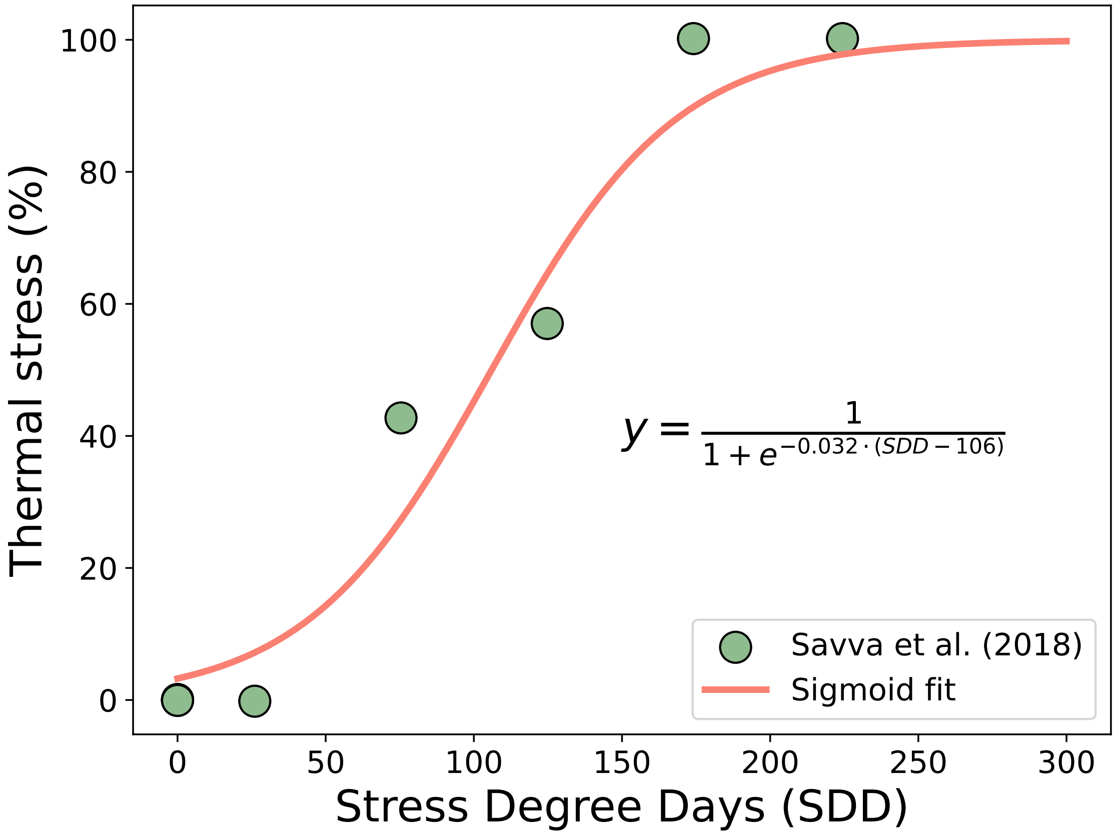


**Supplementary Fig. 4.** Logistic fit relating SDD to observed shoot mortality under constant temperature conditions derived from the experiments from Savva et al. [1].

#### 4 Relationship between $T_{max}$ and seagrass cover and fragmentation

To evaluate how conventional temperature metrics relate to seagrass status, we compared the maximum monthly sea surface temperature ($T_{\max}$) at each site with the estimated *P. oceanica* cover and fragmentation index. Both relationships show significant but moderate correlations, with decreasing cover and increasing fragmentation with $T_{\max}$ (**??**). Notably, substantial degradation in the meadow structure occurred well below the lethal temperature threshold ($LT_{50} = 28.9$ °C) derived from experimental data [1], underscoring that cumulative sublethal exposure, rather than instantaneous lethal exceedance, drives the observed declines.

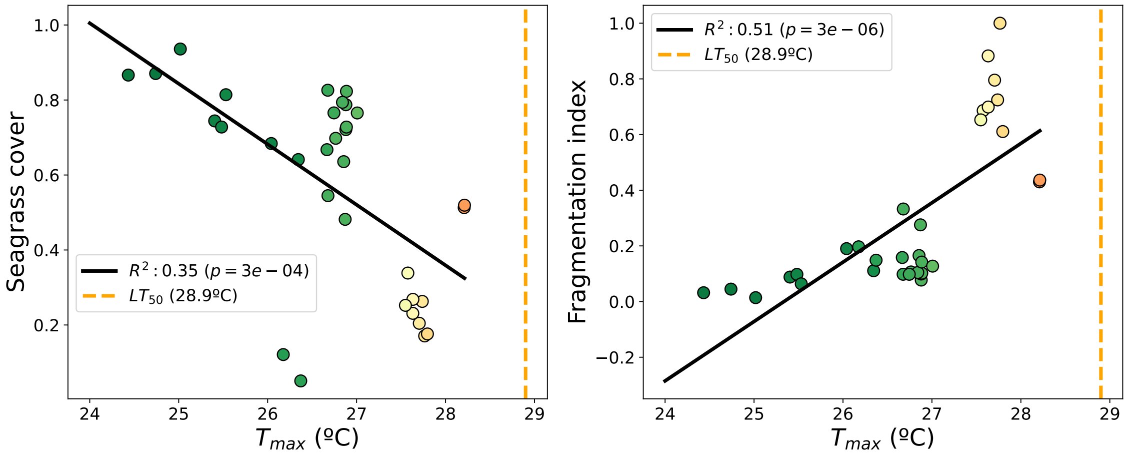


**Supplementary Fig. 5. Relationship between the maximum temperature and seagrass condition.** (a) Relationship between maximum monthly sea surface temperature ($T_{\max}$) and *P. oceanica* cover across Mediterranean sites. (b) Relationship between $T_{\max}$ and the fragmentation index. The dashed orange line indicates the lethal temperature threshold ($LT_{50}$ = 28.9 °C) as reported by Savva et al. (2018) [1]. Both panels show that while mortality-based thresholds capture the upper limit of thermal stress, substantial degradation in meadow condition occurs well below $LT_{50}$.

### 5 Comparison of fragmentation indices

We compared two complementary fragmentation indices to evaluate their consistency and sensitivity to meadow structure: the gap-weighted fragmentation index (FI) and the composite landscape fragmentation index integrating standardized spatial metrics. Both indices are strongly correlated with the number of meadow patches (Supplementary Fig. 6), confirming that they capture similar aspects of structural discontinuity. The high determination coefficients ($R^2 = 0.86$ and $R^2 = 0.77$) indicate robustness across sites and support their joint use in assessing fragmentation patterns in *P. oceanica* meadows.

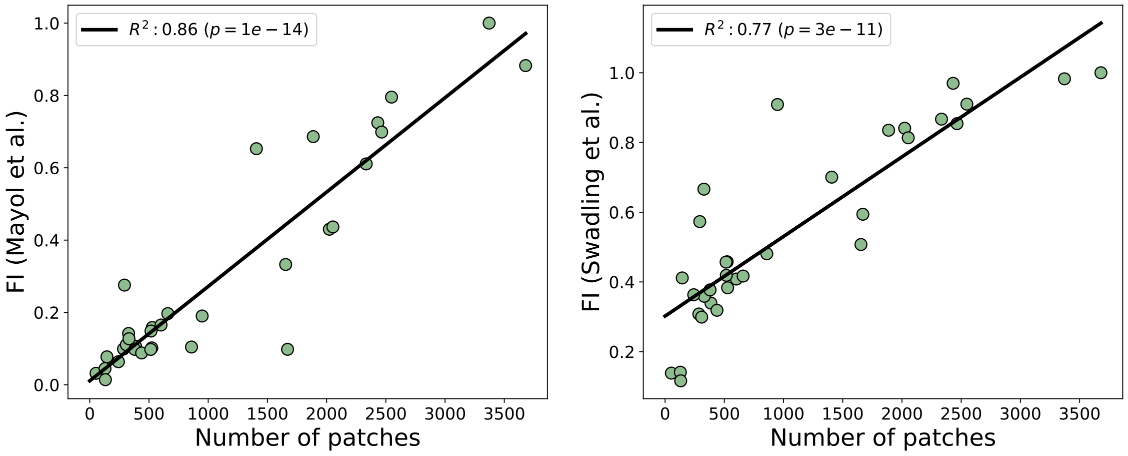


**Supplementary Fig. 6. Comparison between the two fragmentation indices.** Correlation between the gap-weighted fragmentation index (FI, following Mayol et al. 2024 [citation]) and the composite landscape fragmentation index ( Swadling et al. 2023 [2]). Both indices exhibited strong positive correlations with the number of meadow patches, indicating consistent sensitivity to structural discontinuities. Regression statistics ($R^2$ and $p$ values) are shown for each relationship.

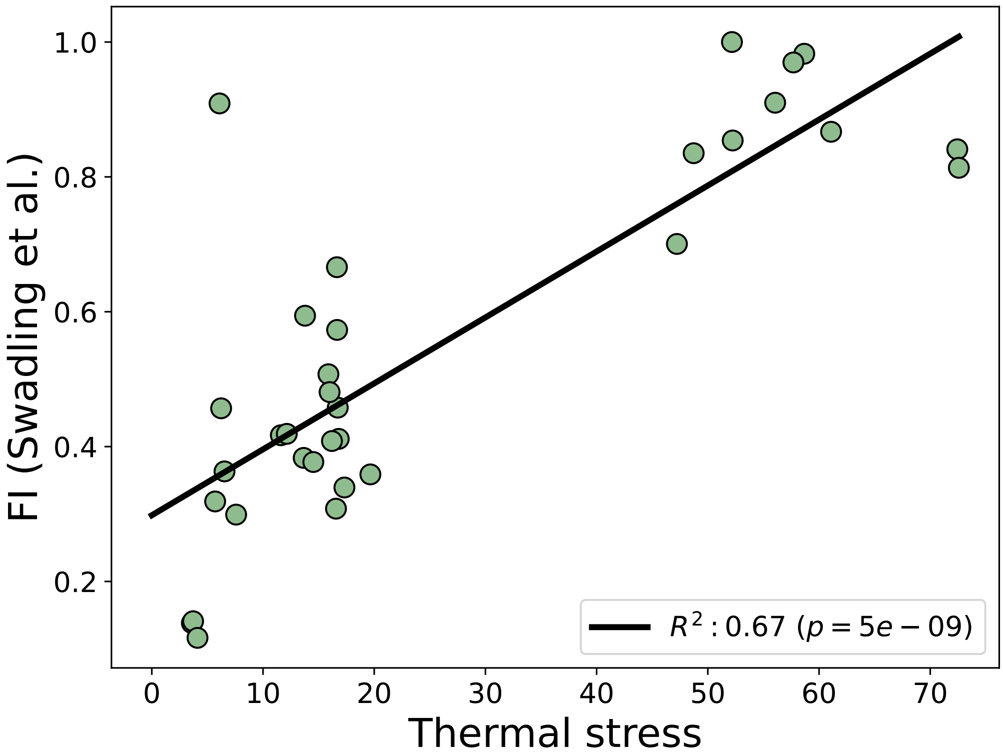


**Supplementary Fig. 7. Relationship between thermal stress and fragmentation.** Scatterplot showing the relationship between cumulative thermal stress (SDD) and composite fragmentation index. Fragmentation increases with thermal stress, supporting the role of cumulative sublethal warming in driving meadow disintegration.

#### 6 Patch size and inter-patch distance distributions

To further characterize the spatial organization, we computed the frequency distributions of patch sizes and inter-patch distances for all study sites. Both distributions exhibit heavy-tailed behavior, consistent with scale-invariant fragmentation (Supplementary Fig. 8). The patch sizes followed a power-law distribution with exponent $\beta = 1.56$, whereas the inter-patch distances exhibited a steeper decay ($\beta = 2.38$). These scaling patterns suggest self-organized spatial clustering and long-range heterogeneity in the *P. oceanica* meadow configurations, reminiscent of the critical transitions observed in other patterned ecosystems.

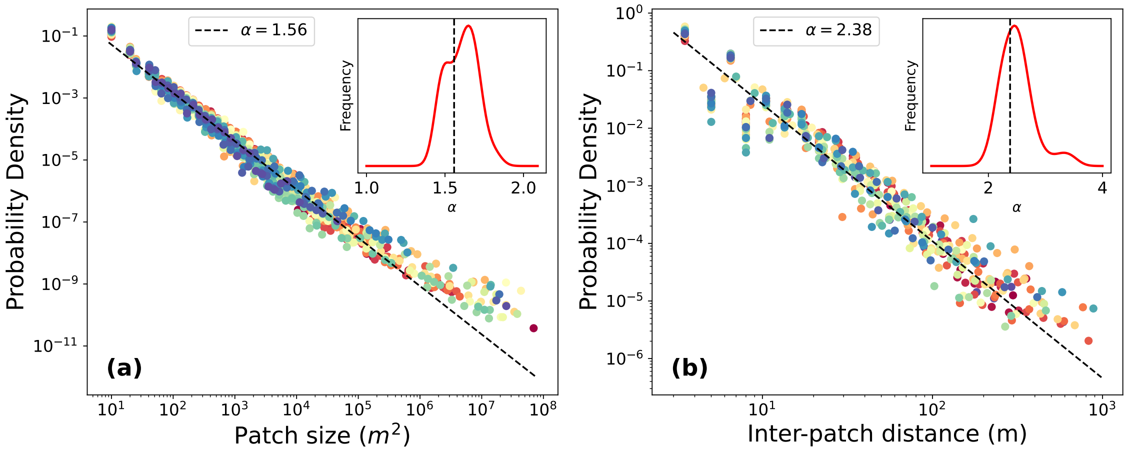


**Supplementary Fig. 8. Patch size and inter-patch distance distributions.** (a) Probability density function of patch size across all study sites, showing a broad, heavy-tailed distribution characteristic of spatial fragmentation processes. The slope of the fitted power-law ($\beta = 1.56$) reflects scale-invariant structure in meadow patchiness. (b) Distribution of inter-patch distances, which also follows an approximate power-law ($\beta = 2.38$), consistent with self-organized spatial clustering in *P. oceanica* meadows.

### 7 Thermal stress under climate change

Future projections of cumulative thermal stress were estimated under RCP4.5 and RCP8.5 scenarios using the Copernicus dataset "Marine biogeochemistry data for the Northwest European Shelf and Mediterranean Sea from 2006 up to 2100 derived from climate projections." Both scenarios predict pronounced intensification and north-eastern expansion of high-stress zones by 2100 (Supplementary Fig. 9). Even under the RCP 4.5 scenario, areas currently exhibiting low to moderate stress are projected to exceed the 50% cumulative stress threshold by 2100, implying widespread degradation and potential contraction of *P. oceanica* meadows across much of the northern Mediterranean. Notably, the distribution of thermal stress is quite heterogeneous, showing the development of new hot spots of thermal stress with climate change, such as the Balearic Islands or south-western Italy. However, some natural refuges were also identified, such as southern Spain, southern France, and the Aegean Sea, which would remain potentially safe even under an RCP 8.5 scenario (Supplementary Fig. 9 h).

To translate these thermal stress projections into structural outcomes, we applied empirically derived relationships between observed SDD and seagrass cover/fragmentation from our 33 Mediterranean sites. The resulting spatial forecasts for seagrass cover, cover change, and fragmentation across the Mediterranean, presented in Supplementary Figs. 10 to 12, paint a stark, diverging picture of *P. oceanica's fate*, contingent on future greenhouse gas emissions. Our results reveal a consistent and worrying pattern: thermal stress does not merely reduce meadow extent but fundamentally compromises its structural integrity. Under the moderate RCP 4.5 scenario, the *P. oceanica* habitat progressively retreats to its thermal refugia in the cooler northern and western coasts (e.g., southern Spain and the Aegean Sea, as identified in Supplementary Fig. 9). By 2099, while significant losses occur in the southern and eastern regions, a substantial portion of the northern Mediterranean still retains moderate to high cover (Supplementary Fig. 10 g), indicating a regionalized decline. In sharp contrast, the high-emissions RCP 8.5 scenario projects a catastrophic regime shift. The initial areas of high-stress expansion—such as the Levantine Sea, the African coast, and the new hotspots around the Balearic Islands and southwestern Italy—quickly become epicenters of habitat collapse. The spatial extent of *P. oceanica* projected for 2099 under RCP8.5 (Supplementary Fig. 10 h) is almost entirely white, signifying the near-total extirpation of functional meadows across the entire basin. This decline is vividly captured in Supplementary Fig. 11, where the dark red color, representing over 80% cover loss, blankets most of the Mediterranean Sea by the end of the century.

Crucially, this loss is intertwined with structural breakdown. Supplementary Fig. 12 illustrates the projected increase in meadow fragmentation, derived from the field-validated correlation between thermal stress and patch disintegration. As early as 2050, the increasing fragmentation (darker orange areas, indicating a higher index value) in the RCP 8.5 scenario precisely matches the expanding thermal hotspots. By 2099, the RCP 8.5 map is almost entirely dark orange, confirming that the cumulative thermal load is expected to transition the few remaining areas of seagrass from continuous, functional meadows into a collection of highly isolated, non-viable fragments. This structural collapse represents an ecological tipping point, rendering the remaining seagrass incapable of sustaining critical ecosystem services, irrespective of the few remaining patches. These projections emphasize that failing to mitigate emissions will rapidly dismantle the foundation species of the Mediterranean coastal ecosystem.

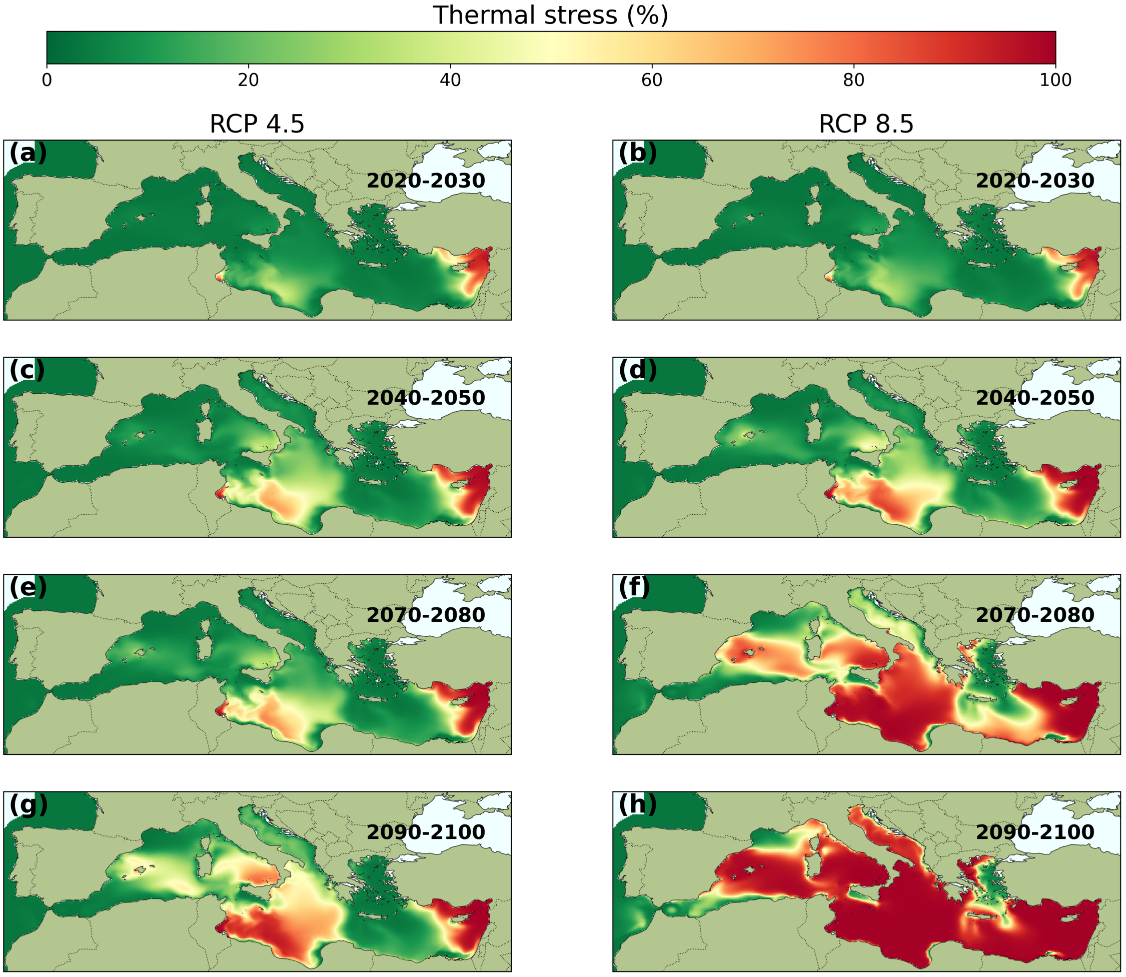


**Supplementary Fig. 9. Projected thermal stress under climate change scenarios.** Spatial distribution of cumulative thermal stress (SDD) in the Mediterranean Sea under future climate projections. Maps correspond to RCP4.5 and RCP8.5 scenarios based on simulations from the Copernicus "Marine Biogeochemistry Data for the Northwest European Shelf and Mediterranean Sea" dataset [3]. The depicted projections correspond to the average results from the indicated time periods. The results highlight a northward expansion of high-stress areas, indicating potential large-scale degradation of *P. oceanica* meadows under continued warming.

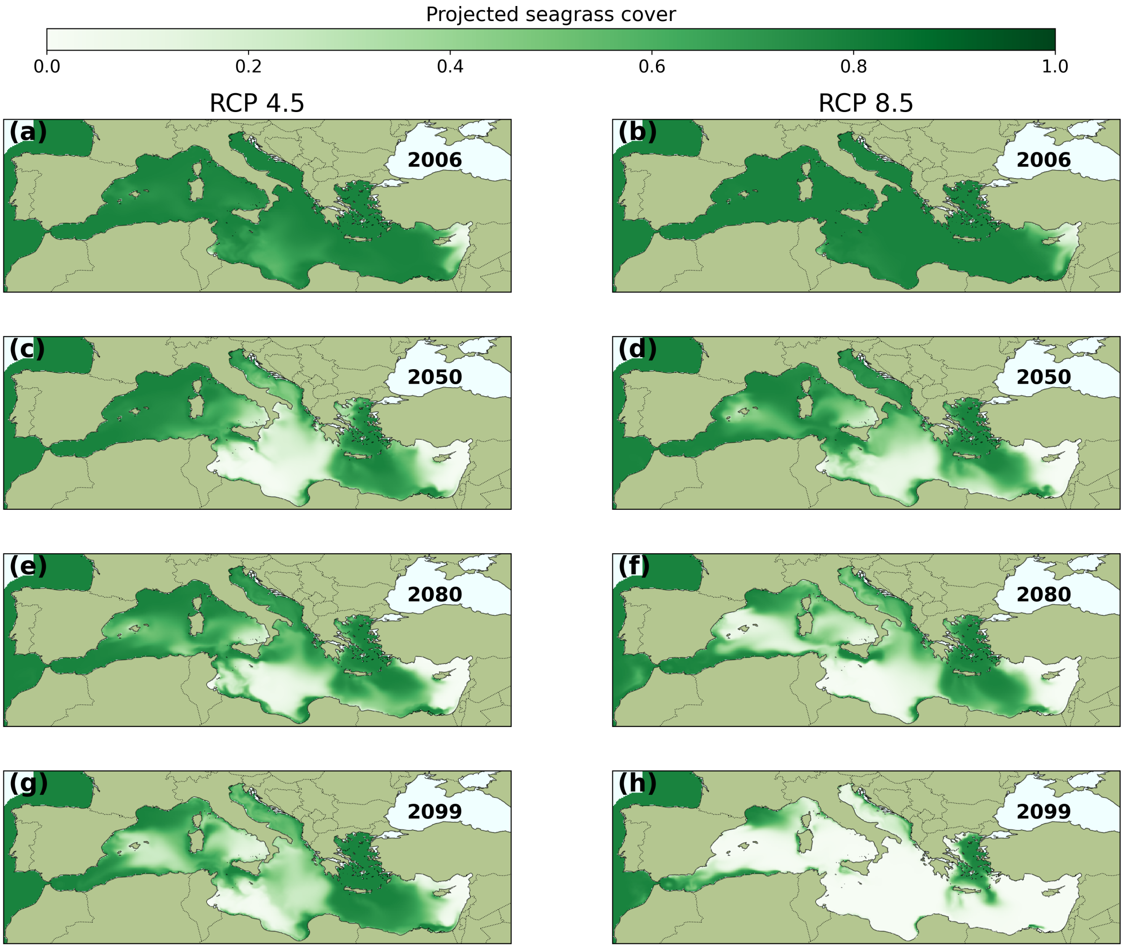


**Supplementary Fig. 10. Projected seagrass cover under climate change scenarios.** Projected *Posidonia oceanica* cover under RCP4.5 (left column) and RCP8.5 (right column) scenarios for 2006, 2050, 2080, and 2099 (rows from top to bottom). Maps show the projected cover derived from the correlation between current SDD and observed seagrass cover across the Mediterranean.

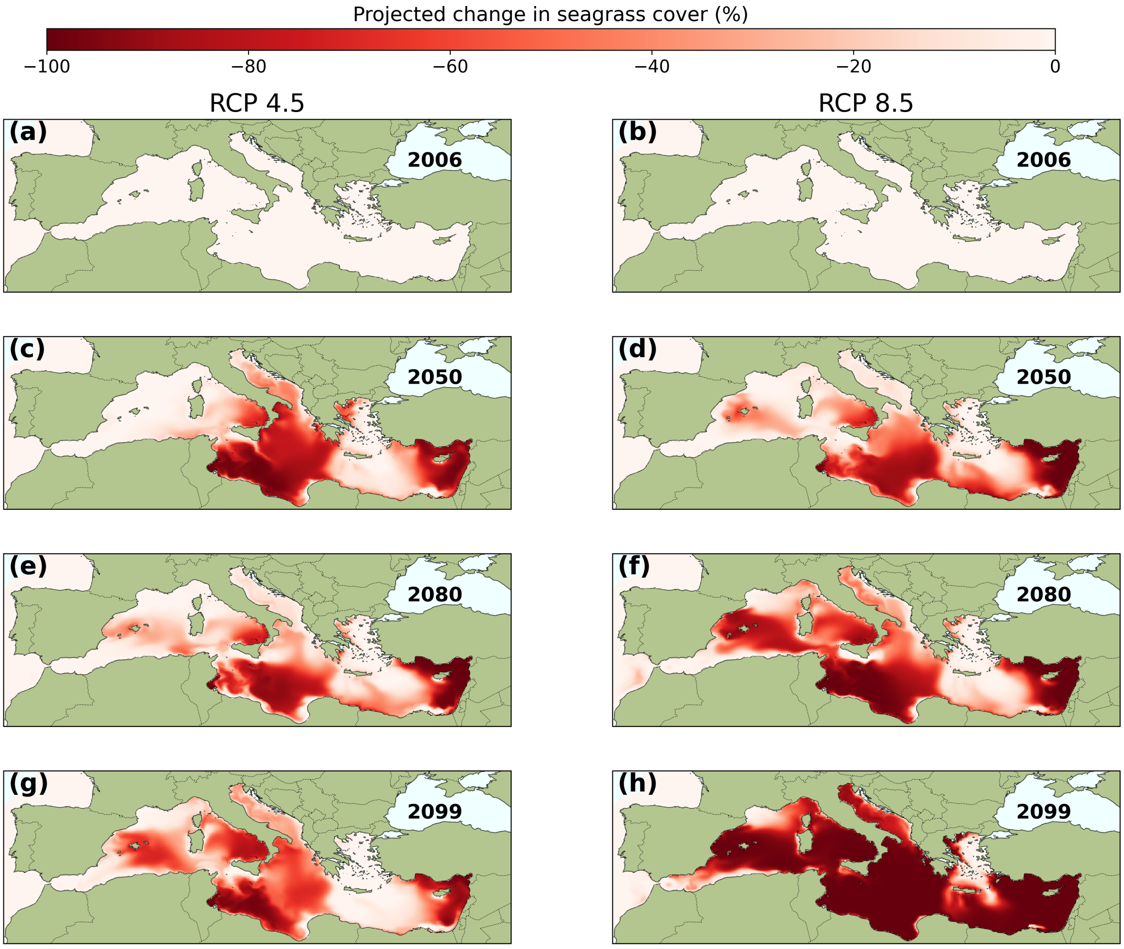


**Supplementary Fig. 11. Projected seagrass cover change under climate change scenarios.** Projected *Posidonia oceanica* cover change under RCP4.5 (left column) and RCP8.5 (right column) scenarios for 2006, 2050, 2080, and 2099 (rows from top to bottom). Maps show the projected change in cover derived from the correlation between current SDD and observed seagrass cover across the Mediterranean.

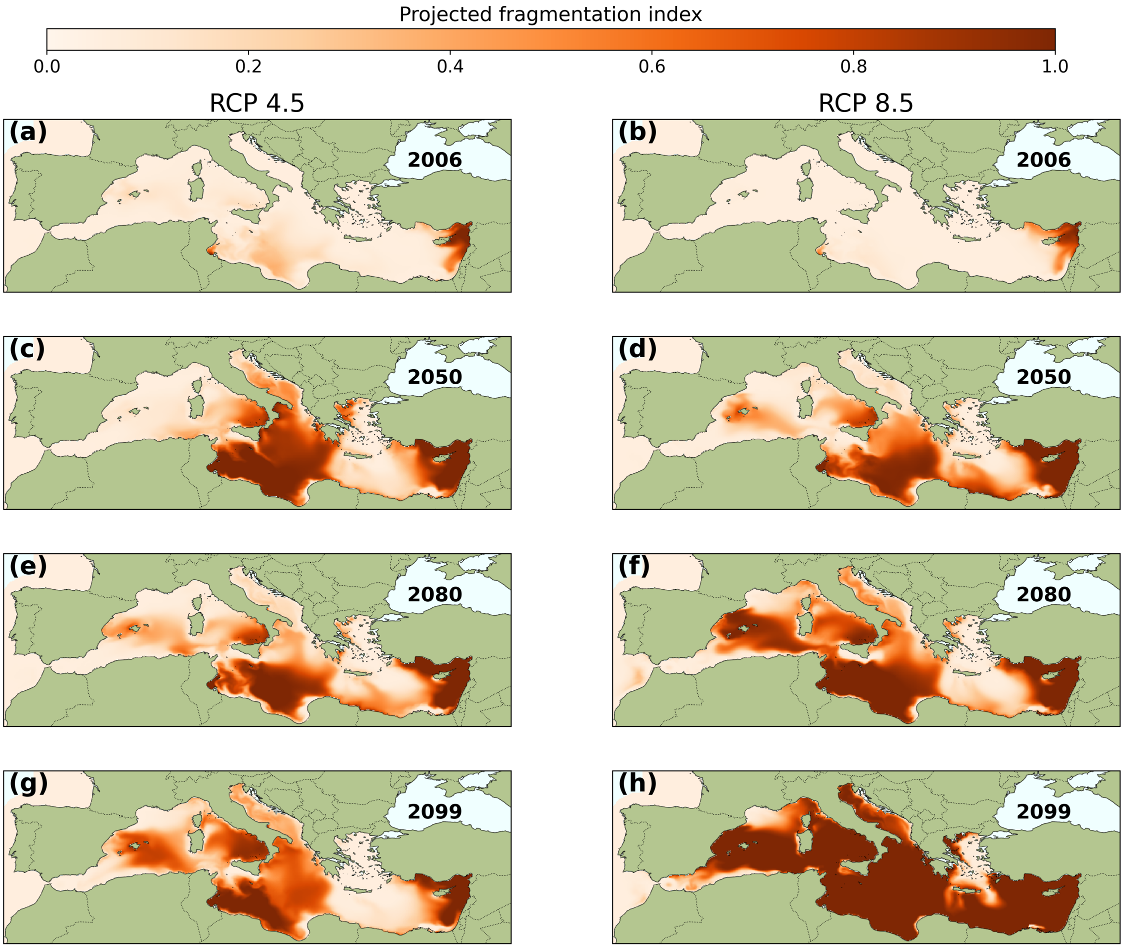


**Supplementary Fig. 12. Projected seagrass fragmentation under climate change scenarios.** Projected *Posidonia oceanica* fragmentation under RCP4.5 (left column) and RCP8.5 (right column) scenarios for 2006, 2050, 2080, and 2099 (rows from top to bottom). Maps show the projected fragmentation derived from the correlation between current SDD and observed seagrass fragmentation across the Mediterranean.
